# LSD impairs working memory, executive functions, and cognitive flexibility, but not risk-based decision making

**DOI:** 10.1101/532234

**Authors:** Thomas Pokorny, Patricia Duerler, Erich Seifritz, Franz X. Vollenweider, Katrin H. Preller

**Affiliations:** Neuropsychopharmacology and Brain Imaging, Department of Psychiatry, Psychotherapy and Psychosomatics, Psychiatric Hospital, University of Zurich, Zurich, Switzerland; Heffter Research Center Zurich, Department of Psychiatry, Psychotherapy and Psychosomatics, Psychiatric Hospital, University of Zurich, Zurich, Switzerland; Department of Psychiatry, Psychotherapy and Psychosomatics, Psychiatric Hospital, University of Zurich, Zurich, Switzerland

## Abstract

Psychiatric and neurodegenerative illnesses are characterized by cognitive impairments, in particular deficits in working memory, decision making, and executive functions including cognitive flexibility. However, the neuropharmacology of these cognitive functions is poorly understood. The serotonin (5-HT) 2A receptor might be a promising candidate for the modulation of cognitive processes. However, pharmacological studies investigating the role of this receptor system in humans are rare. Recent evidence demonstrates that the effects of Lysergic acid diethylamide (LSD) are mediated via agonistic action at the 5-HT_2A_ receptor. Yet, the effects of LSD on specific cognitive domains using standardized neuropsychological test have not been studied. Therefore, we examined the acute effects of LSD (100µg) alone and in combination with the 5-HT_2A_ antagonist ketanserin (40mg) on cognition, employing a double-blind, randomized, placebo-controlled, within-subject design in 25 healthy participants. Executive functions, cognitive flexibility, spatial working memory, and risk-based decision-making were examined by the Intra/Extra-Dimensional shift task (IED), Spatial Working Memory task (SWM), and Cambridge Gambling Task (CGT) of the Cambridge Neuropsychological Test Automated Battery. Compared to placebo, LSD significantly impaired executive functions, cognitive flexibility, and working memory on the IED and SWM, but did not influence quality of decision-making and risk taking on the CGT. Pretreatment with the 5-HT_2A_ antagonist ketanserin normalized all LSD-induced cognitive deficits. The present findings highlight the role of the 5-HT_2A_ receptor system in executive functions and working memory and suggest that specific 5-HT_2A_ antagonists may be relevant for improving cognitive dysfunctions in psychiatric disorders.

## Introduction

Most psychiatric and neurodegenerative illnesses are characterized by cognitive impairments (Claesdotter, *et al*, 2018; Jagust, 2018; Wang, *et al*, 2019; Wunderli, *et al*, 2016). These deficits have a deleterious effect on patients' quality of life and are severely impairing real world functioning (Millan, *et al*, 2012). Previous research has described trans-diagnostic impairments in various cognitive domains in patients (Millan, *et al*, 2012). In particular, deficits in executive functions, working memory, and decision making are among the most common affected domains in psychiatric disorders and have been observed in highly prevalent illnesses like depression, schizophrenia, and substance use disorders (Jessen, *et al*, 2018). While some existing pharmacological treatments have been shown to improve cognitive performance, these effects are small (Désaméricq, *et al*, 2014). Other currently used drugs such as first-generation antipsychotics may even worsen cognitive functions (Hill, *et al*, 2010). Deficits in cognitive abilities are therefore a highly important trans-diagnostic dimension in psychiatric and neurological disorders with great need for improved treatment (Millan, *et al*, 2012).

Pharmacological studies offer the opportunity to causally investigate the contribution of specific receptors and therefore elucidate the neuropharmacological basis of cognitive deficits. This knowledge is urgently needed for the development of specific and novel treatment approaches. Abnormal signalling of the serotonin (5-HT) 2A receptor has been reported in various psychiatric disorders (Zhang and Stackman, 2015). Furthermore, this receptor is widely distributed in brain regions important for cognition and learning (Zhang and Stackman, 2015). However, pharmacological studies investigating the role of this receptor system in humans are rare (Zhang and Stackman, 2015).

Lysergic acid diethylamide (LSD) is a classical hallucinogenic compound and has been shown to transiently induce subjective psychedelic experiences as well as alterations in brain activity and connectivity via agonistic activity on the 5-HT_2A_ receptor (Preller, *et al*, 2018a; Preller, *et al*, 2017; Preller, *et al*, 2018b). The administration of LSD therefore offers the opportunity to causally elucidate the role of the 5-HT_2A_receptor in human cognition. Two recent studies reported that LSD subjectively decreased concentration and increased the self-report of cognitive disorganization (Carhart-Harris, *et al*, 2016; Schmid, *et al*, 2015). Yet, objective measures of cognitive abilities under LSD are still lacking. Therefore, the present study investigated the acute effects of LSD on executive functions, spatial working memory, and risk-based decision-making using computerized and standardized tests provided by the Cambridge Neuropsychological Automated Test Battery (CANTAB). Furthermore, while previous studies point to the involvement of the 5-HT_2A_ receptor in LSD-induced effects (Preller, *et al*, 2017), the contribution of this receptor subtype to alterations in cognitive processes is unknown. To be able to investigate the specific role of the 5-HT_2A_ receptor in cognition, we blocked this receptor subtype via the pre-treatment of LSD with the 5-HT_2A_ receptor antagonist ketanserin. We hypothesized that 1) LSD impairs executive functions, spatial working memory, and risk-based decision-making and 2) that these alterations are attributable to LSD’s agonistic activity on the 5-HT_2A_ receptor.

## Methods

### Participants

Twenty-five healthy participants (19 men, 6 women, mean age ± SD: 25.24±2.79, mean verbal IQ ± SD: 108.4±9.2) were enrolled in the study. All participants underwent a screening procedure at the Department of Psychiatry, Psychotherapy and Psychosomatic, Psychiatric Hospital, University of Zurich consisting of a psychiatric interview (*M.I.N.I., (Sheehan, et al, 1998)*, laboratory test (blood chemistry and urinalysis for drug and pregnancy screening), and a routine medical examination including electrocardiogram. Verbal intelligence was measured with the German version of a multiple choice vocabulary intelligence test (Lehrl, 2005). Volunteers were included when they were 20-40 years of age and willing to refrain from consuming psychoactive drugs at least two weeks before the first experimental session and during the study. The exclusion criteria were personal or first-degree relative history of psychiatric disorders, acute or chronic physical illness, cardiovascular diseases, history of head trauma, neurological diseases such as migraine headaches and epilepsy, history of drug dependence or abuse, a previous significant adverse response to a hallucinogenic drug, and pregnancy or lactation. Before participating, all participants gave their written consent after having received detailed written and oral information about the aims of the study, and the effects and possible risks of the substances administered in accordance with the Declaration of Helsinki. The study was approved by the Ethics Committee of the Department of Public Health of the Canton of Zurich, Switzerland, and the use of LSD was authorized by the Swiss Federal Office for Public Health, Department of Pharmacology and Narcotics, Berne, Switzerland. The current data were collected as part of a larger study (Kraehenmann, *et al*, 2017; Preller, *et al*, 2017; Preller, *et al*, 2018b) and the study was registered at clinicaltrials.gov (NCT02451072).

### Study design and experimental procedures

This study employed a double-blind, randomized, placebo-controlled, within-subject design with three experimental sessions, each separated by at least two weeks. All participants underwent three drug conditions: placebo+placebo (Pla), placebo+LSD (LSD), and ketanserin+LSD (Ket+LSD). One hour after the intake of the first capsule (placebo: 179mg mannitol, 1mg aerosol, p.o.; or ketanserin: 40mg, p.o.) the second one (placebo: 179mg mannitol, 1mg aerosol, p.o.; or LSD: 100µg, p.o.) was administered. All substances were filled in identical looking gelatine capsules. A urine test for drug-screening and pregnancy-test was conducted at the beginning of each experimental session before drug administration. Participants completed the Intra/Extra-Dimensional shift task (IED), Spatial Working Memory task (SWM), and Cambridge Gambling Task (GCT, CANTABeclipse 5.0.12, Cambridge Cognition Ltd., Cambridge, UK) on a computer with a 18″ touch-sensitive screen (Elo Touch Solutions) in a quiet room 220 minutes after the administration of the second capsule. Participants completed the Five Dimension Altered State of Consciousness (5D-ASC) questionnaire (Dittrich, 1998) 720 minutes after drug intake to retrospectively rate subjective drug effects.

### Questionnaire and cognitive tasks

#### Altered States of Consciousness Rating Scale (5D-ASC)

The 5D-ASC (Dittrich, 1998) was used to assess subjective drug effects in each session. Scores were calculated for eleven validated subscales (Studerus, *et al*, 2010): experience of unity, spiritual experience, blissful state, insightfulness, disembodiment, impaired control and cognition, anxiety, complex imagery, elementary imagery, audio-visual synesthesia, and changed meaning of percepts. Results of the 5D-ASC data are expressed as percentage scores of maximum absolute subscale values.

#### Executive Functions: Intra/Extra-Dimensional shift task (IED)

The IED is a test of rule acquisition and cognitive shifting and represents a computerised analogue of the Wisconsin Card Sorting test. Besides attentional set-shifting the task measures different cognitive abilities such as discriminative learning, reversal learning, formation of an attentional set, shifting of attention within the same dimension (intra-dimensional shift, IDS) and between different perceptual dimensions (extra-dimensional shift, EDS). Perceptual dimensions are operationalized via two stimulus characteristics (purple-filled shapes or white lines). Attentional set-shifting is a measure of cognitive flexibility and executive functions. The task consists of nine learning stages, during which the participant first has to focus on shapes or lines within a relevant dimension (IDS) and then shift attention to a previously irrelevant dimension (EDS, stage 8, for a detailed description see Pantelis, *et al*, 2009). The measures of performance on this task are: number of stages completed (stages completed), total number of errors made across all stages adjusted for stages not completed (number of errors adjusted) representing a measure of subject's efficiency in attempting the test, the number of errors on individual stages (number of errors), and the mean time to reach a decision within individual stages (total latency).

#### Working Memory: Spatial Working Memory task (SWM)

The SWM is a self-paced task during which an increasing number of boxes (four, six, and eight) is presented in four trials on the screen. Participants are supposed to find blue tokens which are hidden inside the boxes by touching the boxes and thereby opening them. In each trial the same number of tokens has to be found as the number of boxes presented on the screen. Participants are informed that once a token has been found within a particular box, the box will not be used again to hide a token. This tests measures executive functions (strategy score) as well as working memory errors (between and within errors). Between errors occur when a participant revisited a box in which a token had previously been found, whereas within errors are the number of times a participant revisited a box already found to be empty during the same search sequence. Strategy score describes the use of an efficient search strategy by beginning with a particular box and then returning to that box when a blue token was found to start the new search sequence. High strategy scores represent poor use of a strategy.

#### Risk-based Decision-making: Cambridge Gambling Task (CGT)

The CGT assesses decision-making and risk-taking behaviour outside a learning context. Participants start the task with 100 points. They are presented with 10 boxes on the screen. The boxes are either red or blue. The ration of red and blue boxes varies across trials (9:1, 8:2, 7:3, 6:4, 5:5). Participants have to decide whether a randomly hidden token is more likely to be in a red or blue box. Subsequently, participants bet on their decision by selecting a proportion of their points. The proportion of points (5%, 25%, 50%, 75%, 95%) that can be selected are presented in either ascending or descending order. The outcome measures of this task are: the proportion of trials on which subjects chose the more likely outcome (quality of decision making) and the proportion of current points that the subject stakes on each gamble when the more likely outcome is selected (risk taking).

#### Statistical analysis

Data were analyzed using STATISTICA 8.0 for Windows (StatSoft). For 5D-ASC ratings, a repeated-measures ANOVA with drug (Pla, LSD, Ket+LSD) and subscale (experience of unity, spiritual experience, blissful state, insightfulness, disembodiment, impaired control and cognition, anxiety, complex imagery, elementary imagery, audio-visual synesthesiae, changed meaning of percepts) as within-subject factors were computed. For CANTAB outcome variables (IED: number of errors adjusted, numbers of errors, and total latency; SWM: between errors, within errors, and strategy; CGT: quality of decision making, risk taking) repeated-measures ANOVAs with drug (Pla, LSD, Ket+LSD) as within-subject factor was computed. For the IED stage (1-9) was introduced as additional within-subject factor. For the SWM, stage (4, 6, 8 boxes) was introduced as additional within-subject factor. For the CGT risk ratio (9:1, 8:2, 7:3, 6:4) was introduced as additional within-subject factors. Tukey post-hoc comparisons followed significant main effects or interactions. Pearson’s correlation analyses between CANTAB change scores (drug minus placebo) and 5D-ASC change scores (drug minus placebo) and verbal IQ were computed in case of significant drug effects on CANTAB outcome measures. Statistical comparisons of all data were carried out on a significance level set at p<0.05 (two-tailed).

## Results

### 5D-ASC

There was a significant drug x subscale interaction (F(20,480)=15.10, p<0.000001) (**Fig. 1**), a significant main effect of subscale (F(10,240)=16.62, p<0.000001) and a significant main effect of drug (F(2,48)=85.06, p<0.000001). Tukey post-hoc tests revealed that LSD significantly increased all subscale scores compared to Pla and Ket+LSD (all p<0.0001) except for anxiety (p>0.7). There were no significant differences between Pla and Ket+LSD in any subscale score (all p>0.9)

**Figure 1:**
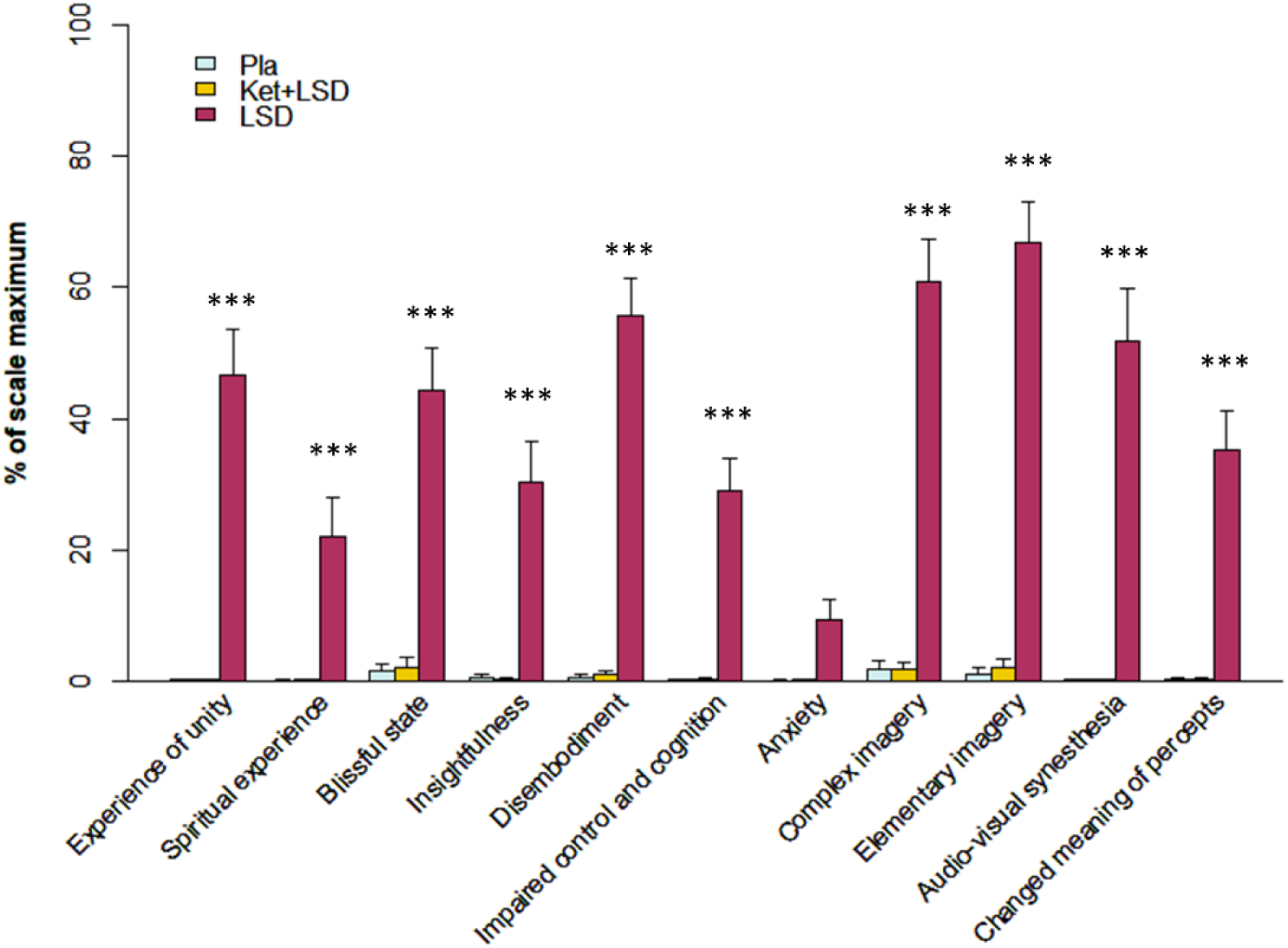
Subjective drug effects assessed with the 5D-ASC questionnaire 720 min after substance administration. LSD significantly increased all scale scores compared to Pla and Ket+LSD (all p<0.0001), except for anxiety (p>0.7). Data are expressed as mean + SEM. ***p<0.0001, corrected.

### Intra/Extra-Dimensional shift task

The stages completed did not differ between drug conditions (F(2,48)=1.5, p>0.2; mean (SD) Pla: 8.84 (0.11), Ket+LSD: 8.76 (0.13), LSD: 8.68 (0.15)). However, for number of errors adjusted, there was a significant effect of drug (F(2,48)=4.9, p<0.05)) with more errors in the LSD condition compared to Pla (p<0.05) and Ket+LSD (p<0.05) (**Fig 2A**). To further investigate at which stage the errors occurred, we computed a repeated-measures ANOVA with stage and drug as within-subject factors. We found a significant drug x stage interaction (F(16,384=1.81, p<0.05) (**Fig. 2B**), a significant main effect of stage (F(8,192)=9.10, p<0.00001), and a significant main effect of drug (F(2,48)=4.50, p<0.05). Tukey post-hoc tests revealed that LSD significantly increased the number of errors in stage 8 (EDS) compared to Pla (p<0.0001) and Ket+LSD (p<0.01), but not in any other stage compared to both, Pla and Ket+LSD (all p>0.9). There were no significant differences in the number of errors in any stage between Pla and Ket+LSD (p>0.9). For total latency, there was a significant drug x stage interaction (F(16,384)=1.94, p<0.05) (**Fig. 2C**), a significant main effect of stage (F(8,192)=12.47, p<0.00001), and a significant main effect of drug (F(2,48)=7.46, p<0.01). Tukey post-hoc tests revealed that LSD significantly increased latency in stage 8 (EDS) compared to Pla (p<0.0001) and Ket+LSD (p<0.0001), but not in any other stage compared to Pla (p>0.6) or Ket+LSD (p>0.7). There were no significant differences in total latency scores in any stage between Pla and Ket+LSD (all p>0.9).

**Figure 2:**
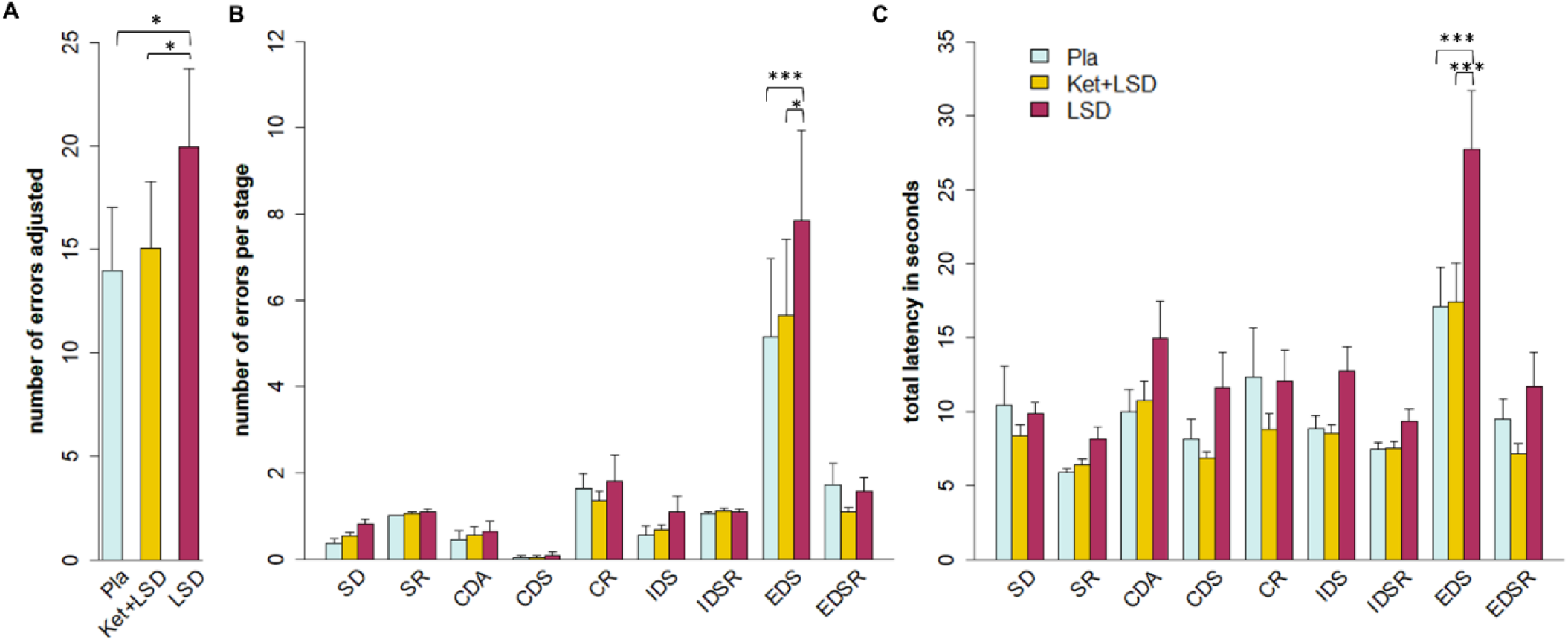
Intra/Extra-Dimensional shift task. A) displays the total number of errors adjusted for stages completed. LSD significantly increased the number of errors adjusted compared to Pla and Ket+LSD (both p<0.05, corrected). B) displays the number of errors in each stage. LSD significantly increased the number of errors in the EDS stage compared to Pla (p<0.0001, corrected) and Ket+LSD (p<0.01). C) displays the total latency (mean time to reach a decision within individual stages). LSD significantly increased latency in the EDS stage compared to Pla and Ket+LSD (both p<0.0001). Data are expressed as mean + SEM. *p<0.01, ***p<0.0001. IED stages: *SD* simple discrimination, *SR* simple reversal, *CDA* compound discrimination adjacent, *CDS* compound discrimination superimposed, *CR* compound reversal, *IDS* intra-dimensional shift, *IDSR* intra-dimensional shift reversal, *EDS* extra-dimensional shift, *EDSR* extra-dimensional shift reversal

### Spatial Working Memory

For between errors, there was a significant drug x stage interaction (F(4,96)=4.59, p<0.01) (**Fig. 3A**), a significant main effect of drug (F(2,48)=7.14, p<0.01), and a significant main effect of stage (F(2,48)=34.88, p<0.00001). Tukey post-hoc tests revealed that participants made significantly more between errors in the LSD condition than in the Pla condition when six boxes were presented (p<0.01). Further, LSD significantly increased between errors when eight boxes were presented compared to both Pla (p<0.001) and Ket+LSD (p<0.001). There were no significant differences in the number of between errors between Pla and Ket+LSD at any stage (all p>0.6). For within errors, there was no significant drug x stage interaction (F(4,96)=1.00, p>0.4) or main effect of drug (F(2,48)=1.84, p>0.1) (**Fig. 3B**). However, we found a significant main effect of stage (F(2,48)=5.09, p<0.01). For the strategy score, there was a significant drug x stage interaction (F(4,96)=4.11, p<0.01) (**Fig. 3C**), a significant main effect of stage (F(2,48)=205.50, p<0.00001), but no significant main effect of drug (F(2,48)=1.48, p>0.2). Tukey post-hoc tests revealed that the strategy score was increased (reflecting poor use of a strategy) under LSD compared to Pla and Ket+LSD when eight boxes were presented (both p<0.01). There was no significant difference in the strategy scores between Pla and Ket+LSD at any stage (all p>0.9).

**Figure 3:**
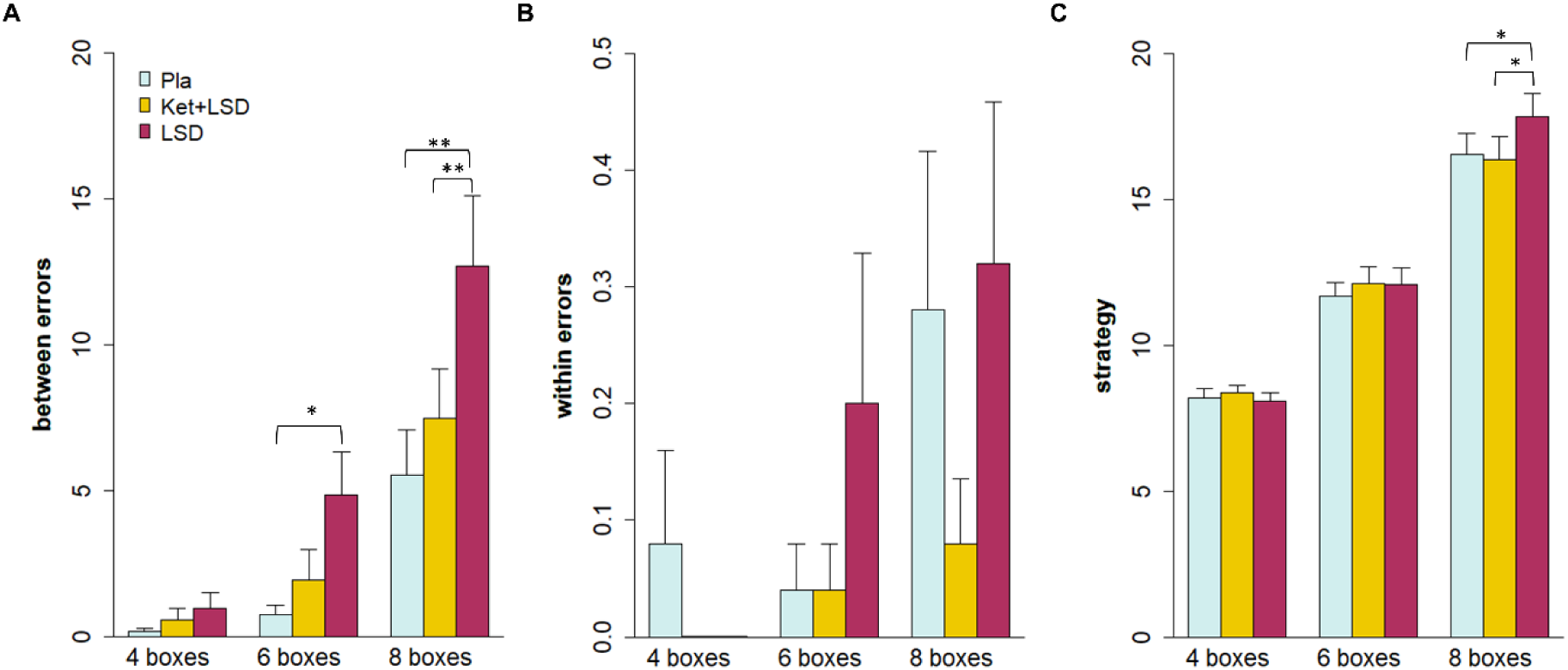
Spatial Working Memory. A) displays between errors. LSD significantly increased the number of between errors when six boxes were presented compared to Pla (p<0.01, corrected). When eight boxes were presented, LSD significantly increased between errors compared to Pla and Ket+LSD (both p<0.001, corrected). B) displays within errors. There were no significant drug effects for number of within errors. C) displays the strategy scores. LSD reduced the use of an efficient search strategy when eight boxes were presented compared to Pla and Ket+LSD (both p<0.01). Data are expressed as mean + SEM. *p<0.01, **p<0.001.

### Cambridge Gambling Task

For quality of decision making, there was no significant drug x risk ratio (F(6,144)=0.38, p>0.8), interaction (**Fig. 4A**), and no main effects of drug (F(2,48)=0.50, p>0.6) or risk ratio (F(3,72)=1.93, p>0.1). For risk taking (**Fig. 4B**), there was no significant drug x risk ratio (F(6,144)=1.04, p>0.4) interaction, and no significant main effect for drug F(2,48)=1.05, p>0.3). However, there was a significant main effect of risk ratio (F(3,72)=108.06, p<0.000001) for risk taking, indicating that participants made higher bets when the risk ratio was lower.

**Figure 4:**
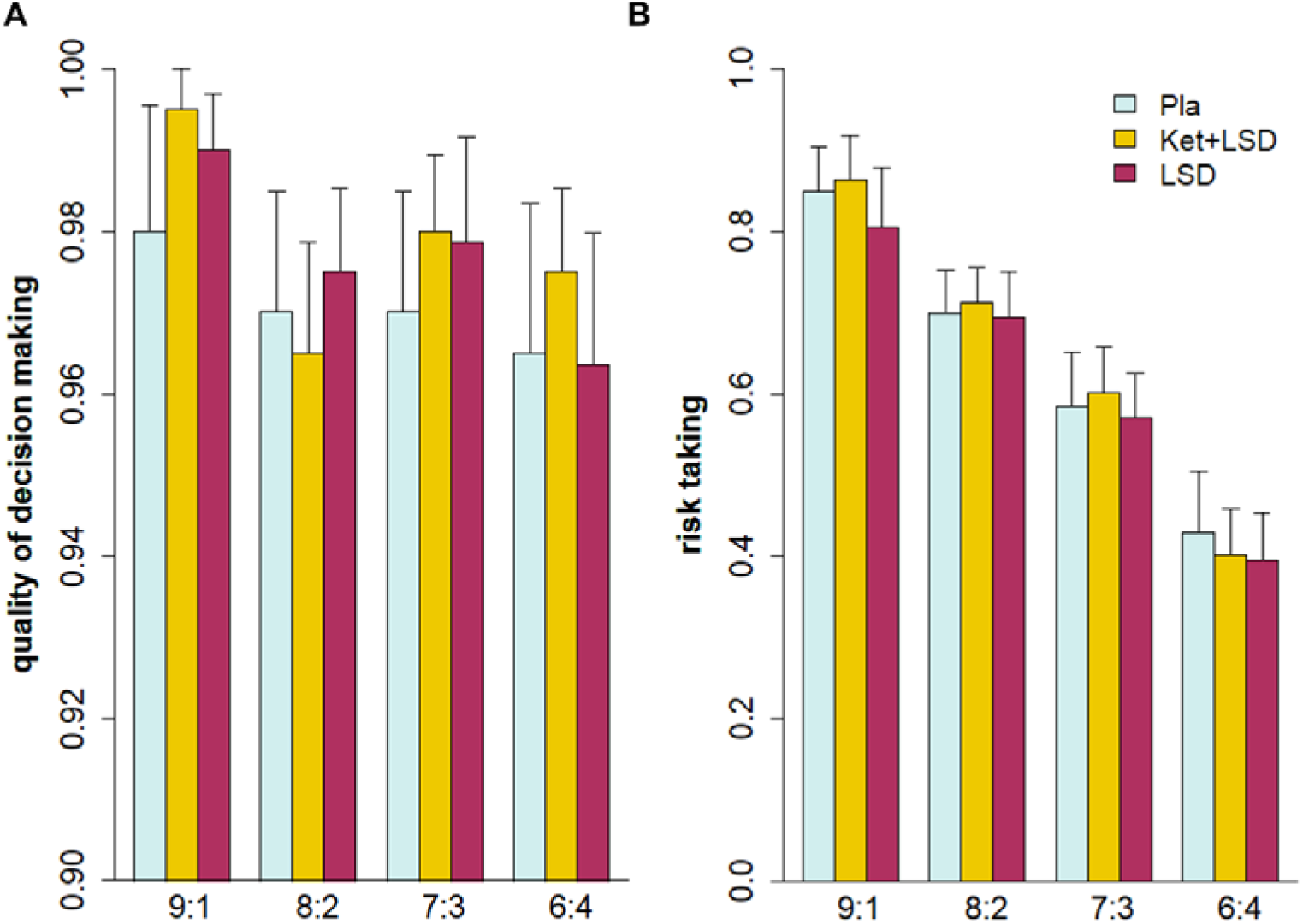
Cambridge Gambling Task. No significant effects of drug were found for A) quality of decision-making or B) risk taking. Data are expressed as mean + SEM.

### Correlations

After Bonferroni correction for multiple comparisons there were no significant correlations between change scores of CANTAB outcome measures and change scores of 5D-ASC subscale scores (all p>0.1, corrected). Furthermore, IQ was not correlated with change scores of CANTAB outcome measures (all p>0.3, corrected).

## Discussion

Trans-diagnostic deficits in cognitive abilities are highly prevalent in psychiatric and neurological disorders, but are insufficiently improved by current treatment approaches (Millan, *et al*, 2012). The current study closes major knowledge gaps in the field via the administration of LSD together with a 5-HT_2A_ receptor antagonist and the application of standardized and computerized cognitive tasks that capture the most relevant cognitive domains impaired in psychiatric disorders (Jessen, *et al*, 2018). We show that I) acutely administered LSD significantly impaired executive functions and working memory compared to placebo, II) risk-based decision-making was unaffected by LSD, and III) LSD-induced cognitive deficits and subjective symptoms were dependent on 5-HT_2A_ receptor stimulation.

### LSD impairs cognitive flexibility and executive functions on the Intra/Extra-Dimensional shift task

LSD led to a significant increase in error rates and increased latency in the EDS stage of the IED task compared to placebo. These impairments were normalized in the Ket+LSD condition. Impairments in the EDS stage are interpreted as signs of preservation of a previously established attentional set (Elliott, *et al*, 1995). Extradimensional shifting requires being able to inhibit the previously established attentional set and shift attention between stimulus dimensions. Therefore, impairments in EDS are interpreted as reduced cognitive flexibility (Chamberlain, *et al*, 2007). Processes that may underlie this LSD-induced decrease in cognitive flexibility are deficits in executive functions, in particular increased susceptibility to distraction from task-irrelevant stimuli (Jazbec, *et al*, 2007). This interpretation is supported by previous studies in healthy humans which have shown that psilocybin, a structurally related serotonergic hallucinogen and 5-HT_2A_ receptor agonist, led to a significant decline of correct detection and an increase of false alarm in the AX continuous performance test (Umbricht, *et al*, 2003) and to a significant reduction in attentional tracking ability on a multiple-object tracking task, an effect attenuated by ketanserin (Carter, *et al*, 2007). The authors of the later study suggest that the impaired attentional performance under psilocybin may reflect a reduced ability to suppress or ignore distracting stimuli rather than reduced attentional capacity per se. This interpretation may also explain seemingly contradictory findings reporting increases in psychological flexibility under the influence of other psychedelic substances (Kuypers, *et al*, 2016). Here, participants were instructed to provide alternative interpretations of the presented stimulus material. However, the applied tasks did not require the suppression of distracting elements which may explain the differing results. In contrast to our current finding, an early study with LSD found no significant change in performance on the Wisconsin Card Sorting Test (Primac, *et al*, 1957), of which the IED is the computerised analogue. It seems likely that this previous study was not able to detect LSD-induced effects due to the small sample size (n=10). Importantly, LSD-induced impairments in cognitive flexibility and executive functions were normalized by ketanserin, therefore pinpointing the crucial role of the 5-HT_2A_ receptor in these cognitive functions.

### LSD impairs executive functions and working memory on the Spatial Working Memory task

LSD compared to placebo led to a significant increase of between errors and decreased use of strategy in the SWM. An increase in between errors represents a deficit in working memory, since participants revisited boxes even though a token had already been found within the box. Deficits in strategy represent impairments in executive functions. Importantly, these effects were only present when cognitive load was high. These results are in line with previous studies investigating the effects of other psychedelic 5-HT_2A_ agonists on spatial working memory. Psilocybin has been shown to dose-dependently impair performance on a spatial working memory task (Wittmann, *et al*, 2007). Furthermore, ayahuasca acutely impaired working memory (Bouso, *et al*, 2013). Similarly to results obtained on the IED, pre-treatment with ketanserin normalized LSD-induced working memory and executive function deficits on the SWM in the current study.

### The role of the 5-HT_2A_ receptor in LSD-induced cognitive impairments

On both, the IED and the SWM task, blocking the 5-HT_2A_ receptor with ketanserin prevented LSD-induced cognitive deficits, therefore pointing to the importance of this receptor system in working memory and executive functions. This result is in line with a recent computational study indicating that 5-HT_2A_ receptors contribute to spatial working memory tasks (Cano-Colino, *et al*, 2014). The hippocampus is particularly involved in spatial memory and has moderate to high levels of 5-HT_2A_ receptors (Dwivedi and Pandey, 1998). Injections of the 5-HT_2A_ receptor antagonist ritanserin in rat hippocampus significantly improved spatial memory in the Morris Water Maze task (Naghdi and Harooni, 2005). Furthermore, the prefrontal cortex (PFC) is implicated in higher-order executive tasks such as working memory, attention, and executive function (Millan, *et al*, 2012). The PFC is linked to the parietal cortex, which exerts a modulatory influence on attention and working memory (Millan, *et al*, 2012). Both structures have a particular high density of 5-HT_2A_ receptors, and they exert top-down modulatory influence on subcortical regions including the hippocampus (Pompeiano, *et al*, 1994). Clinical evidence corroborates the current results pinpointing the importance of the 5-HT_2A_ receptor in cognitive abilities: Atypical antipsychotics, which have 5-HT_2A_ receptor antagonistic properties, have been shown to be advantageous for treating cognitive impairments in schizophrenia compared to classic antipsychotics (Meltzer, *et al*, 2012). There are qualitative similarities between hallucinogen-induced alterations in information processing and the symptoms of an early phase of schizophrenic psychoses (Geyer and Vollenweider, 2008). Specifically, sensorimotor gating as indexed by prepulse inhibition (PPI) is impaired in patients with schizophrenia and related to cognitive deficits (Braff, *et al*, 2001). LSD and psilocybin disrupt PPI in healthy subjects, an effect that was also correlated with impairments of sustained attention in healthy humans (Gouzoulis-Mayfrank, *et al*, 1998; Quednow, *et al*, 2012; Schmid, *et al*, 2015; Vollenweider, *et al*, 2007). Importantly, it has been shown that the disruptions of PPI induced by psilocybin in humans and LSD in rats are reversed by 5-HT_2A_ receptor antagonists (Halberstadt and Geyer, 2010; Ouagazzal, *et al*, 2001; Quednow, *et al*, 2012). Together these results suggest that the 5-HT_2A_ receptor system may be a promising target in the treatment of trans-diagnostic impairments in working memory and executive functions.

### LSD does not affect risk-based decision making on the Cambridge GamblingTask

Interestingly, in the CGT, no significant differences between the three drug conditions were found. LSD did not influence the quality of decision making and risk taking. Decision making and risk taking may therefore represent cognitive domains that are not modulated by 5-HT_2A_ receptor signalling. This is in line with a recent study that showed that psilocybin had no effect on moral decision-making (Pokorny et al., 2017). Furthermore, this result supports previous reports that associate risk-based decision-making behaviour with dopaminergic signalling, whereas the serotonin system has been suggested to play a role in the regulation of cognitive biases, and therefore the appraisal of reinforcers when selecting between actions, in particular in a learning context (Rogers, 2011). The CGT, however, is not relying on learning, and risk-based decision making on the CGT may therefore not be sensitive to alterations in 5-HT_2A_ receptor signalling. Yet, LSD also has affinity for dopamine receptors and animal studies have reported a first 5-HT_2A_ receptor mediated and a second D2 receptor mediated phase of action (Marona-Lewicka and Nichols, 2007; Marona-Lewicka, *et al*, 2005). However, the involvement of the dopaminergic system in this study is unlikely as pretreament with ketanserin normalized the LSD-induced effects not only in the CANTAB tasks 220 minutes after drug intake but also the retrospective rated psychological effects measured with the 5D-ASC. Therefore, the present results indicate that the effects of LSD in humans are primarily mediated via 5-HT_2A_ receptor activation. This is in line with recent reports, that LSD in humans increased the levels of prolactin and cortisol, which are markers for serotonergic drug activity (Schmid, et al, 2015).

However, it is possible that higher doses of LSD are needed to modulate dopaminergic activity and potentially induce alterations in risk-based decision making.

### Correlations and limitations

Cognitive impairments in the IED and SWM did not correlate with the LSD-induced subjective effects. Importantly, this suggests that cognitive processes under LSD are not confounded by psychedelic effects, in particular visual inaccuracies or disturbances. Furthermore, LSD-induced impairments were not related to individuals’ IQ, suggesting that 5-HT_2A_ receptor stimulation by LSD impaired working memory and executive functions independently of general intelligence.

Previous studies have shown that the selective 5-HT_2A_ receptor antagonist ketanserin is suitable for studying the role of 5-HT_2A_ receptors in human performance and to investigate the specific contribution of the 5-HT_2A_ receptor to effects of psychedelic drugs (Carter, *et al*, 2007; Kometer, *et al*, 2012; Liechti, *et al*, 2000; Quednow, *et al*, 2012; Vollenweider, *et al*, 1998; Preller, *et al*, 2017). A limitation of the present study is the lack of a fourth drug condition investigating the effect of ketanserin alone. However, previous studies have shown that ketanserin neither led to any significant differences in subjective drug effects assessed with the 5D-ASC (Carter, *et al*, 2007; Kometer, *et al*, 2012), nor to performance changes in cognitive tasks such as a spatial working memory task (Carter, *et al*, 2007), or the Stroop task (Quednow, *et al*, 2012).

## Conclusion

In conclusion, the present study pinpoints the role of the 5-HT_2A_ receptor in cognitive processes, in particular executive functions, cognitive flexibility, and spatial working memory. However, risk-based decision-making outside a learning context was unaffected by LSD and is therefore potentially not mediated by the 5-HT_2A_ receptor. Blocking the 5-HT_2A_ receptor by ketanserin normalized both LSD-induced cognitive impairments and subjective drug effects. As altered 5-HT_2A_ receptor density and cognitive dysfunctions are found in several psychiatric disorders such as in schizophrenia, autism, and obsessive compulsive disorder (Millan, *et al*, 2012), this receptor subtype represents a promising target to help understanding the neurophramacological basis of cognitive processes and to improve treatment in affected patients.

## Acknowledgments

We thank Amanda Planzer, Jan Flemming, and Rainer Krähenmann for their assistance in data collection.

